# Genome sequence assembly evaluation using long-range sequencing data

**DOI:** 10.1101/2022.05.10.491304

**Authors:** Dengfeng Guan, Shane A. McCarthy, Jonathan M. D. Wood, Ying Sims, William Chow, Zemin Ning, Kerstin Howe, Guohua Wang, Yadong Wang, Richard Durbin

## Abstract

Genome sequences are computationally assembled from millions of much shorter sequencing reads. Although this process can be impressively accurate with long reads, it is still subject to a variety of types of errors, including large structural misassembly errors in addition to localised base pair substitutions. Recent advances in long single molecule sequencing in combination with other long-range technologies such as synthetic long read clouds and Hi-C have dramatically increased the contiguity of assembly. This makes it all the more important to be able to validate the structural integrity of the chromosomal scale assemblies now being generated. Here we describe a novel assembly evaluation tool, Asset, which evaluates the consistency of a proposed genome assembly with multiple primary long-range data sets, identifying both supported regions and putative structural misassemblies. We present tests on three *de novo* assemblies from a human, a goat and a fish species, demonstrating that Asset can identify structural misassemblies accurately by combining regionally supported evidence from long read and other raw sequencing data. Not only can Asset be used to assess overall assembly confidence, and discover specific problematic regions for downstream genome curation, a process that leads to improvement in genome quality, but it can also provide feedback to automated assembly pipelines.

## Background

*De novo* assembly is essential to generate reference genome sequences for new species. Until recently this has relied on computational assembly of millions of short fragmented DNA sequences from next generation sequencing (NGS) machines. The assembly obtained in this way is often known to be problematic [1] due to repetitive sequences, sequencing bias, heterozygous alleles, etc. Repeat sequences that make up a major proportion of all eukaryotic species genomes [2], can trap the assembly algorithm and lead to misassemblies, especially when the repeat size is larger than the read size, which is typical for NGS data. Sequence composition bias, also often called GC bias, and can further affect genome completeness when portions of the genome have very high or low GC content [3].

In the last few years, the advent of long read sequencing technologies, such as PacBio Single Molecule Real-Time (SMRT) sequencing and Oxford Nanopore Technology sequencing (ONT), which have read lengths at least two orders of magnitude larger than NGS data and exhibit much less sequencing bias, has been revolutionising genome assembly studies. This has stimulated the development of new genome assembly tools such as Falcon [4], which is a string graph-based assembler[5], and Canu [6], an overlap graph-based assembly tool. The application of long reads in genome assembly has greatly improved assembly continuity and completeness compared to that of NGS assembly [7].

Ambitious new *de novo* sequencing projects are taking advantage of these long read technologies, such as the Vertebrate Genomes Project (VGP) which hopes to generate *de novo* assemblies for all the vertebrate species [8] and ultimately the Earth BioGenome Project, which advocates sequencing over 1.5 million eukaryotic species, including animals, plants and microbiomes, in the next ten years [9]. With the number of *de novo* assemblies likely to be released over the coming years a method is needed to reliably evaluate their accuracy.

A number of metrics are frequently used to assess a draft assembly. In particular, genome continuity is typically measured by contig or scaffold N50, which is the longest length such that at least half of the total sequence is in contigs or scaffolds longer than this. The larger the N50, the more continuous the assembly is, and so researchers prefer tools that can generate a larger N50. However N50 can only reflect the continuity of the assembly: one can force erroneous joins to make a larger N50, and in this case the assembly quality is poor even with a large N50. Another assembly metric, genome completeness, can be gauged by BUSCO [10], which relies on single-copy orthologs to quantify genome completeness.

However, neither of the above metrics measure the accuracy of the draft assembly. Several tools have been developed to address this. GAGE (Genome Assembly Golden-standard Evaluation) [11] compares a set of NGS data assemblies based on deep sequencing data and summarises putative misassemblies into two categories, indels, and misjoins which is subclassified into inversions, relocations (within chromosomes) and translocations (between chromosomes). QUAST [12] and QUAST-LG [13] require a reference, with respect to which they give a detailed description of a proposed assembly, such as NA50 which is the N50 of the aligned assembly blocks, and again the numbers of misassemblies split into relocations, inversions and translocations. They also supply various visualized results for better understanding the assembly quality.

However, these tools rely on an existing reference genome to find misassemblies, which makes them not applicable to newly sequenced species. Even if a reference genome exists, the natural mutations between the samples may still lead to false positive misassemblies. To address this, tools such as Amosvalidate [14], REAPR [15] and Tigment [16] all evaluate a *de novo* assembly against primary data sets. Amosvalidate applies a method of combining multiple sources of misassembly signals, including mate-pair orientation, repeat content, read depth, micro-heterogeneities, and read alignment breakpoints, to detect suspicious misassembled regions. However, it was developed before the invention of NGS technologies and requires an AMOS Bank format as input, which are not generated by most of the genome assemblers. REAPR uses independent paired-end reads alignments to calculate a fragment coverage distribution (FCD) for each base and use FCD outliers to pinpoint misassemblies, and Tigment calculates molecular coverage for the whole assembly from linked reads, and then uses that as evidence for misassembly detection.

Although all these tools can be used to detect misassemblies, none of them uses the full collection of long-range sequencing data that is now typically available, which in principle should provide more reliable results. We therefore designed and developed a tool “Asset” which uses the four types of long-range sequencing datasets currently used by the VGP, namely PacBio long reads, 10X linked reads, Bionano optical maps, and Hi-C [17], to identify suspect misassemblies. We performed experiments on three *de novo* assemblies, and demonstrated that none of the data types individually are sufficient to validate a genome assembly, but that in combination “Asset” can identify misassemblies accurately. With more and more *de novo* sequencing projects being launched, Asset has the potential to be applied systematically to help accelerate and standardise the genome evaluation process. Furthermore, it can provide lists of potential problems for subesequent genome curation to focus on [18, 19], and rank genome assemblers.

## Results

### Asset methods overview

As described above, Asset is able to use information from four primary data types. We first briefly describe these and the principles behind the information that they provide.

Because they sequence single molecules without amplification subject to composition bias, current long read platforms give rather uniform sampling across the genome. Therefore evenness of long read coverage is one good indicator for identifying misassemblies, where regions with extremely low or high read depth are potentially misassembled. However, this metric can be confused by perfect or near perfect repetitive sequence regions. The reads mapped to these regions are usually ignored for read depth calculation due to ambiguous mapping positions, or alternatively are randomly assigned, both of which can result in anomalous depth and a misassembled region recognized by some tools. Further, sufficiently long repetitive regions will have no spanning reads, and thus are often discarded or treated as misassemblies. Of course, it is precisely around these long repetitive regions that missassemblies are likely to occur.

These problems can be partially solved by using linked reads. Linked-read or “read cloud” data consists of barcoded reads from a long fragment of DNA, such as are produced by 10X Genomics or similar technology. Following alignment to a reference or assembly it is possible to reconstruct the likely extent of the underlying 20-200kb DNA fragments [20], and from this calculate the physical long fragment coverage for each base. Places where the molecule coverage drops to zero or near zero frequently indicate missassemblies. However an exception to this can occur when missing a large piece of sequence over 200kb.

The restriction map is another useful data type for misassembly identification. Although a full nucleotide sequence is not provided, the longer molecule lengths used by modern optical restriction mapping technologies such as BioNano are capable of spanning regions over 250kb and can cover longer repetitive regions. Bionano’s new direct label and stain (DLS) technique, which eliminates repeated breakpoints in the fragments, can give information on very long range structure across tens of megabases. However, there remain some long regions, typically over 1Mb such as centromeres, which are not spanned by BioNano maps, but across which order and orientation can be confirmed using Hi-C reads.

Hi-C is based on Chromatin Conformation Capture technology, where DNA is cut and re-ligated *in situ* in the nucleus, and then following selective purification of chimeric ligation products a short read library is made for deep sequencing [17, 21]. This gives contact information between DNA fragments at multiple resolutions up to chromosome scale, allowing us in principle to use Hi-C data to confirm whether a contig is correctly concatenated with its neighbours regardless of the length of the missing piece.

It is worth noting that the goal of Asset is to confirm that the sequence given is fully supported by the data, not that it is complete in the sense of explaining all the data. This means that even for a diploid assembly which describes both chromosomes, or provides a primary assembly plus alternate “haplotig” material [4], we are able to just assess the primary assembly. This is what we do throughout the rest of this paper.

### Asset pipeline

The Asset pipeline is illustrated in Additional file 1: Figure S1, and described fully in the Methods section. Here we give a brief overview.

Given an input of a primary assembly and four types of long-range data including long reads, linked reads, restriction maps and Hi-C reads, we apply the following strategy to identify suspect misassemblies.

To begin we partition the assembly scaffolds into *sequences,* which are composed of A,C,G,T bases, and *joins,* which are runs of Ns between contigs. We then apply different strategies to confirm sequences and joins.

For the sequences, we align the long reads and restriction maps to the scaffolds. Only considering the primary alignments, so each part of a read/map is aligned in at most one place, and ignoring the 300bp at the end of each read alignment, we label segments of the contigs that are neither covered by long reads nor consistent restriction map intervals as *unsupported sequence*.

To confirm joins we consider three possible types of evidence. First, a restriction map alignment that spans across a join with consistent spacing. Second, for linked reads, we identify molecule extents as intervals containing a sufficient density of mapped reads with the same barcode. We require a sufficient number of inferred molecules to cross a join to confirm it. Third, for Hi-C data, we split all contigs into two equal halves, and calculate the number of Hi-C read pairs linking each pair of contig-halves; to be confirmed, we require that the right contig-half is the best partner of the left contig-half and vice versa, where by “best” we mean that it is the contig-half linked by the greatest normalised number of read pairs. Joins not supported by any of these evidence types are labeled as *unsupported joins*.

Finally, we combine the two sets of unsupported regions, merging regions within 10kb of each other. We also drop isolated regions of unsupported sequence that are within a fixed distance (default: 1kb) of a gap since they are mostly called by natural drop of long read depth. We regard the remaining unsupported regions as suspect misassemblies.

### Evaluation of results

Three different assemblies were evaluated using a variety of primary data, as listed in Table 1.

**Table 1.**
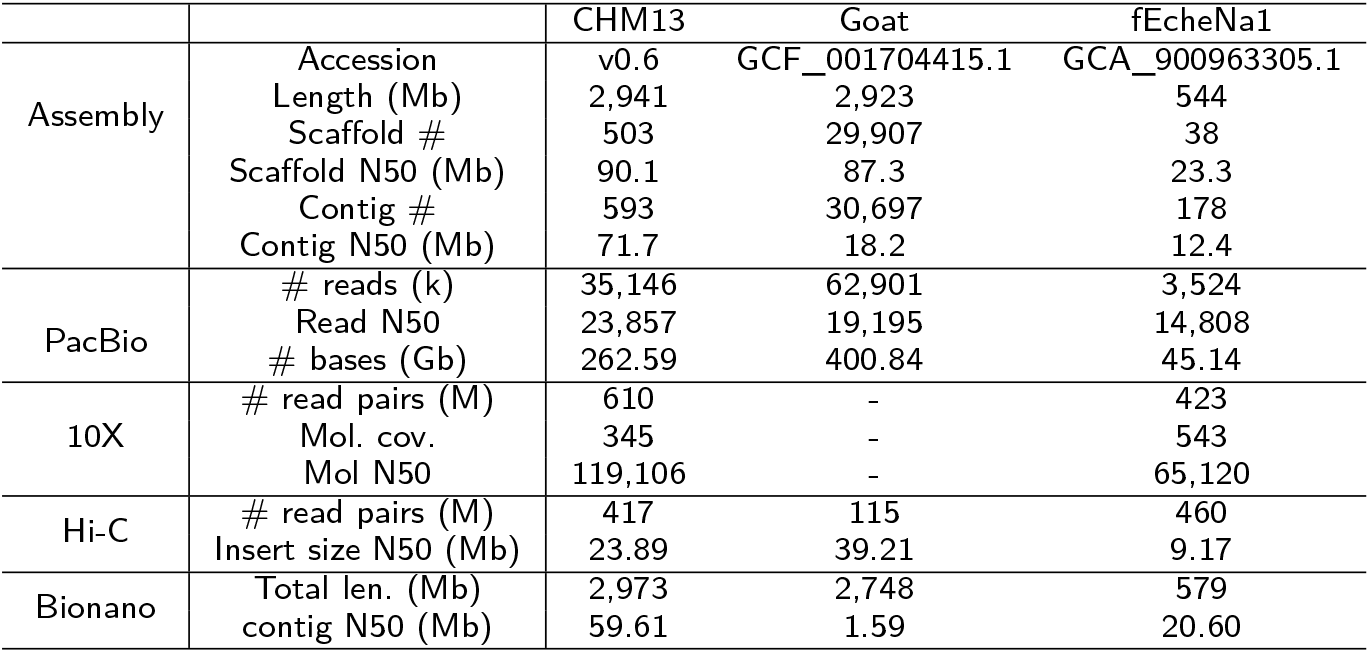
Assemblies and sequencing datasets used for Asset analysis.

To quantify the accuracy for each tool, we examined a representative subset of suspect misassemblies by hand. For Asset results we selected all those on one chromosome, while for the other methods we randomly selected 20 candidates, since they produce too many misassemblies to validate all of them on a chromosome.

To further investigate the types of misassemblies found by Asset, we classified these into nine types: HAPDUP (haplotypic duplication, where both divergent haplotypes are included), HAPMIX (mixed haplotype, where a mosaic of divergent haplotypes is given), DUPSEQ (local sequence duplication), INS (sequence insertion), MIS (sequence deletion), COLL (local sequence collapsing), INV (inversion), RELOC (relocation), TRAN (translocation).

### Application on the CHM13 assembly

We carried out our first experiment on the CHM13 human assembly assembled using Canu v1.7.1 with 39X rel1 Oxford Nanopore data and 70X PacBio reads at scaffold level by the Telomere-to-Telomere (T2T) consortium [22]. This represents the genome of the CHM13hTERT cell line, which was obtained from a hydatidiform mole and hence is homozygous diploid, avoiding problems of heterozygosity. The greatest effort was made on ChrX, where the gaps were manually checked and filled, which created a full length of ChrX. The whole assembly contains 2.94 Gb bases, with scaffold N50 90.1 Mb, and contig N50 71.7 Mb (Table 1).

In this experiment, we used 35 million PacBio reads from 215 runs, 610 million 10X linked read pairs and 417 million Hi-C read pairs. The Bionano consensus map is created from the DLE-1 direct labeling enzyme data (Table 1). We ran QUAST-LG with default settings using the GRCh38 assembly as a reference and REAPR with default settings using the linked reads.

**Table 2.** Misassembly identification results.

|  | Asset | QUAST | REAPR |
| --- | --- | --- | --- |
| CHM13 | 368 | 2,942 | 98,501 |
| Goat_ARS1 | 541 | - | 128,262 |
| fEcheNa1 | 171 | - | 28,304 |

QUAST could only be applied to the human assembly because it requires a reference genome to identify misassemblies, which is not independently available for goat and fEcheNa1.

**Table 3.**
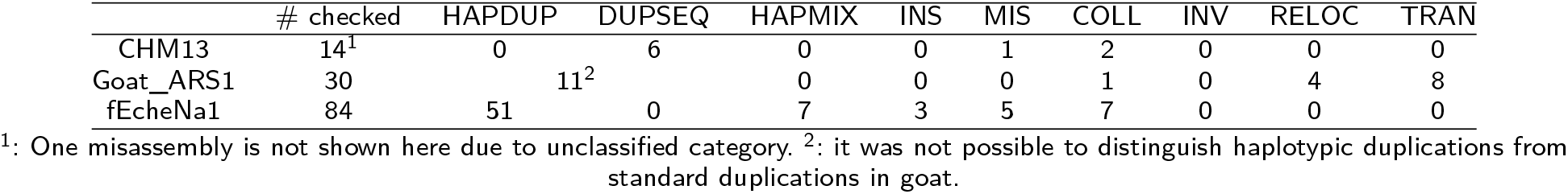
Manually checked Asset misassemblies results.

In our experiment, Asset found 368 misassemblies on the CHM13 super scaffolds. We manually checked 14 misassemblies on two major scaffolds Super-Scaffold_434 (SS434) and Super-Scaffold_445 (SS445) of Chr1, for which we validated 5 and 7 misassemblies respectively. We also validated another 2 candidate misassemblies on the complete ChrX sequence. Of these 14 candidate misassemblies, 10 were confirmed as real, giving a specificity of approximately 71.4%.

QUAST-LG discovered 2,942 extensive misassemblies on the super scaffolds, 20 were randomly picked up for manual validation from the same scaffolds as above, 3 of them appeared to be correctly identified, while yields an accuracy of 15%. QUAST-LG found 345 misassemblies on the complete ChrX assembly, including 319 relocations, 13 translocations and 13 inversions, most of these are recognized due to segmental duplication, centromeres and telomeres which are not well resolved in the GRCh38 assembly. Additional file 1: Figure S2 demonstrates an example of QUAST-LG translocations which are falling into a segmental duplicated region on ChrX and ChrY in the GRCh38 assembly. As is shown in the figure, the start of ChrX is mapped to ChrX and ChrY in the reference assembly (Figure S2a), and QUAST-LG found 8 translocations in that region, which are not true based on the Bionano Access view (Figure S2b). REAPR identified 98,501 misassemblies in total, 20 were randomly chosen for validation, 1 is correct, which results in an accuracy of 5%, around 60% misrecognised misassemblies are caused by repeat elements and the others are misclassified because of uneven read depth distribution.

In the 14 manually checked misassemblies from Asset, we classified them into the nine categories mentioned above, 6 of them are DUPSEQs, 1 is a MIS and 2 are COLLs, 4 are close to the centromere which are hard to classify, one can either be a MIS or INS, which proves CHM13 may not be a complete haploid. We give an example of a DUPSEQ found on SS445 in Figure 1. From the dotplot (Figure 1a), SS445 is mapped to the GRCh38 Chr1 assembly, both two regions ~130-250kb and ~250-370kb on SS445 are mapped to ~700-810kb on NC_000001.11, the duplicate region is marked in red, the PacBio coverage in its corresponding region drops almost to zero. From the Bionano Access (Figure 1b), we can see a clear divergence between the DLE consensus maps and the SS445, which indicates a ~120kb insertion in the scaffold. Combining the evidence from Bionano Access and the dotplot, there is a ~120kb DUPSEQ on the SS445.

**Figure 1.**
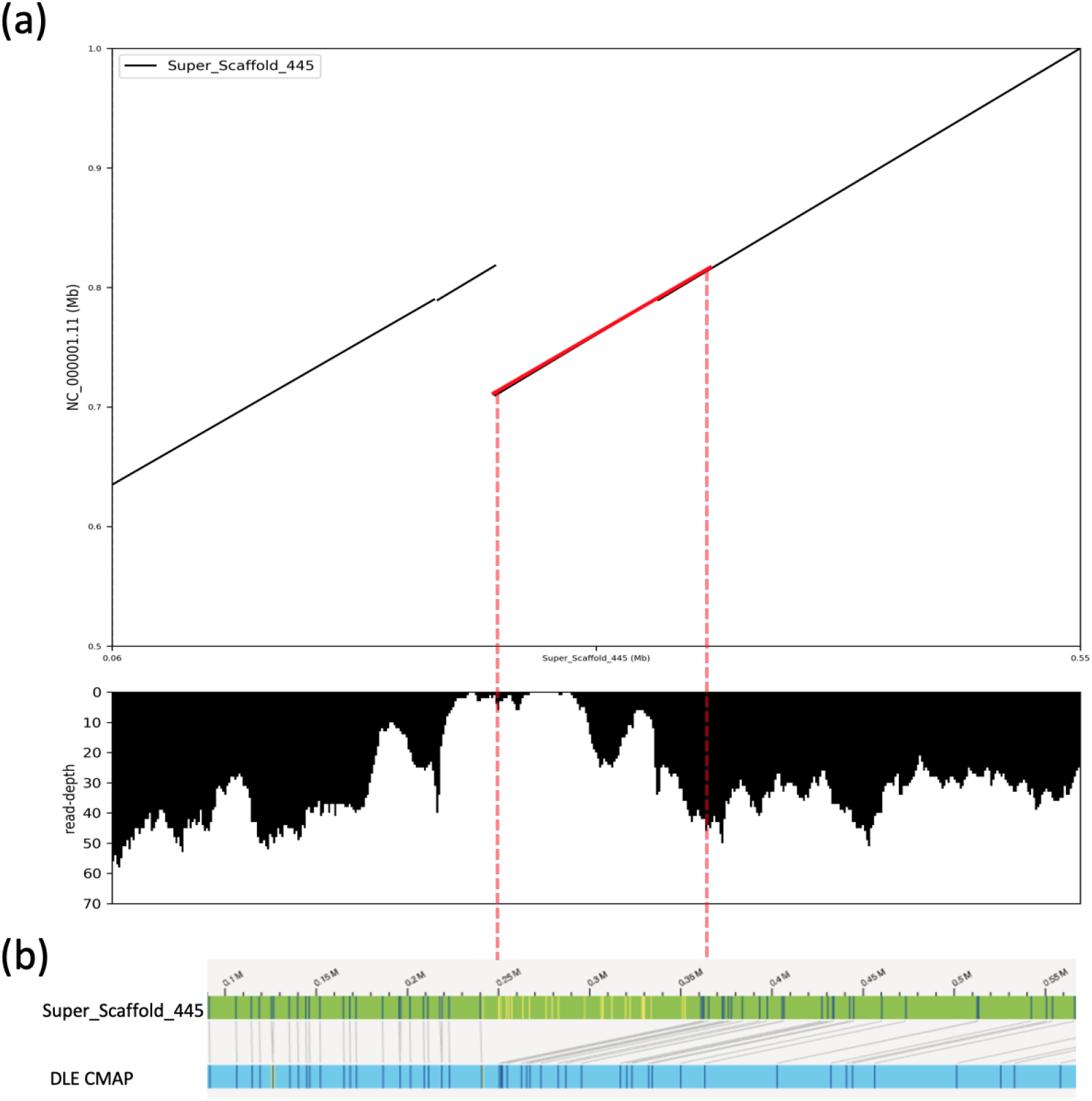
A DUPSEQ on CHM13 SS445. (**a**) dotplot of SS445 mapped to GRCh38 Chr1 (NC_000001.11). The duplicate sequence is marked in red. The PacBio coverage in the duplicated region almost drops to zero. (**b**) Bionano Access view of the alignment between SS445 and the DLE consensus map. Based on alignment, there is a around 120kb large insertion from 0.24 to 0.36 Mb in the original scaffold.

The most intriguing case found by Asset on SS434 may indicate the CHM13hTERT cell is not a comprehensive haploid. The misassembly is located on 12,121,611-12,126,513 of SS434, where the PacBio coverage is low and Bionano alignment is screwed up. Figure 2 illustrates the alignments between the Bionano DLE consensus map and the scaffold for this region. Through comparing the two Bionano consensus maps, we can observe an apparent divergence between them, there is a ~82kb difference between “A-a” and “B-a” regions, where A, B, a are all restriction enzyme sites. Even though the assembler seemed to recover the larger haplotype, it still missed a ~26kb sequence.

**Figure 2.**
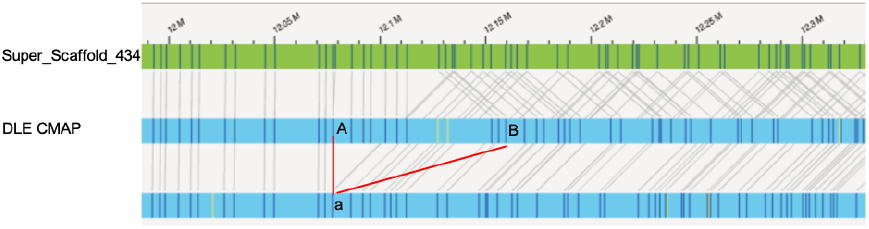
Bionano Access view of CHM13 heterozygous haplotypes on SS434. A, B and a are all restriction enzyme recognition sites. In two consensus maps, both site A and site B are mapped to site a, which indicates an apparent divergence between two haplotypes of SS434. And the region is found because the assembler fails in representing either of the haplotypes.

### Application on the goat reference assembly

The third experiment is applied on a *Capra hircus* assembly (domestic goat, RefSeq assembly accession: GCF_001704415.1) [23]. It was assembled using The Celera Assembler PacBio Corrected Reads pipeline[7] with 69X PacBio subreads, and scaffolded with 98X optical mapping data and 115 million Hi-C read pairs. The final assembly contains 2.92 Gb bases in total, the scaffold N50 is 87.3 Mb, and the contig N50 is 18.2 Mb (Table 1).

In this experiment, Asset used 63 million PacBio reads in 325 runs, the datasets contain a total 400.84 Gb bases, and 115 million Hi-C read pairs (Table 1). Lacking publicly available linked reads for the assembly, we did not use linked read data in the experiment. For Bionano data, we used the Irys BspQI consensus map assembled by the Bionano Access software package and the sequencing data mentioned in the assembly paper is from a male offspring of the original species. REAPR used public Illumina datasets containing 443 million read pairs for the test and ran in default mode.

Asset found 541 misassemblies on chromosomes. We manually checked the 30 misassemblies on Chr1, found 24 were real misassemblies, this gives an approximate accuracy of 80%. We discovered 8 of them were involved in TRANs, 11 were DUPSEQs or HAPDUPs which are hard to classify due to insufficient evidence, 4 were RELOCs and 1 was COLL. REAPR reported 128,262 assembly errors. Among 20 randomly selected misassemblies, 2 were real, which results in an accuracy of 10%.

We chose one asset-identified relocation region on 40.87-41.89Mb of Chr1 and demonstrated it in Figure 3, this region is also identified in the Goat assembly paper. In the HiGlass [24] view (Figure 3a), we can observe a strong signal between the start of the scaffold and this region, indicating this region should be moved to the front of the scaffold. The same situation shows in the dotplot (Figure 3b), where we mapped Chr1 to another assembly CHIR_2.0 (Yunnan Black Goat, GenBank assembly accession GCA_000317765.2) [25], the 40.87-41.89Mb on Chr1 is mapped to the start of the Chr1 of the black goat assembly. Meanwhile, from the Access view (Figure 3c), we can see that two breaks at 40.8Mb and 41.9Mb, the middle regions from 40.8-41.9Mb are aligned continuously, however it is not joined with either side. Based on these evidence, the region found by Asset is a relocation.

**Figure 3.**
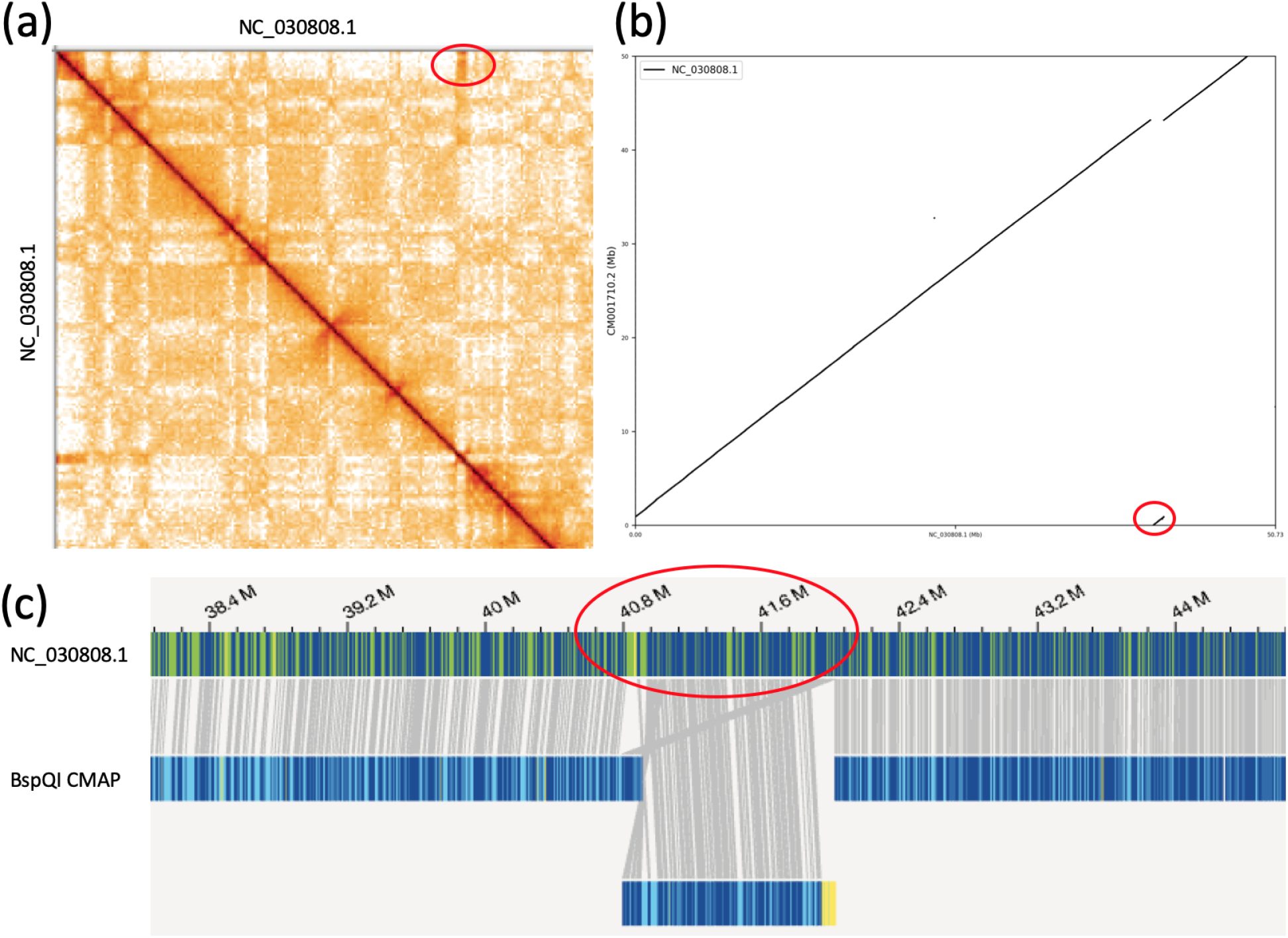
A RELOC on goat Chr1 (NC_030808.1) assembly. (**a**) HiGlass view on Chr1. A strong signal implies the between 0 to 40.8Mb. (**b**) dotplot of 0-50Mb on Chr1 mapped to CM001710.2 (Chr1 of Yunnan black goat assembly). The dotplot indicates that 40.8–41.9Mb of Chr1 should be moved to the start of the chromosome. (**c**) Access view shows the same situation, the middle region is misplaced.

### Application on the VGP fEcheNa1 assembly

Our final experiment is performed on a VGP assembly, *Echeneis naucrates* (fEcheNa1, GenBank assembly accession: GCA_900963305.1) is assembled by Sanger VGP assembly group. It goes through a VGP assembly routine v1.5, where the assembly is built with ~60X PacBio sequencing data using Falcon, uses Falcon-unzip to construct pseudohaplotypes, and scaffolded with 10X reads by using scaffold10x (https://github.com/wtsi-hpag/Scaff10X) and Bionano optical mapping data. The scaffolds finally reached chromosome level with Hi-C data using SALSA [26], and went into a stringent curation process by Sanger GRIT team. The curated assembly contains 38 scaffolds, including a total 544Mb bases, the scaffold N50 is 23.3Mb and contig N50 is 12.4Mb.

With regard to the number of misassembly, the fEcheNa1 assembly is better than the other two tested assemblies, but it still has a few misassemblies. Asset identified a total of 171 misassemblies on chromosomes. We manually check the 84 misassemblies on the scaffolds which are larger than scaffold N50, 73 of them are real which generates an accuracy of 86.9%. Majorities of these misassemblies are HAPDUPs(51, 60.7%), 7 were COLLs, 7 were HAPMIXs, 3 were INSs, 5 were MISs, and the remaining ones locating at the telomeres were hard to classify. REAPR found 28,304 chromosomal misassemblies, 20 were randomly picked up for manual checking, 6 were real misassemblies, which yields an accuracy of 30%.

Figure 4 gives an example of a HAPDUP on Chr13 (NC_042523.1), in the figure, the dotplot illustrates the self versus self alignment of NC_042523.1, we can see the read depth in the middle region goes down to about half of its normal coverage (~60X), and from the Access view, there is a ~170kb from 24.94-25.11Mb, which is corresponding to the duplication. HAPDUP is a common issue for heterozygous assemblies, and the problem is partially resolved by a tool “purge_dups”[27] developed by the authors.

**Figure 4.**
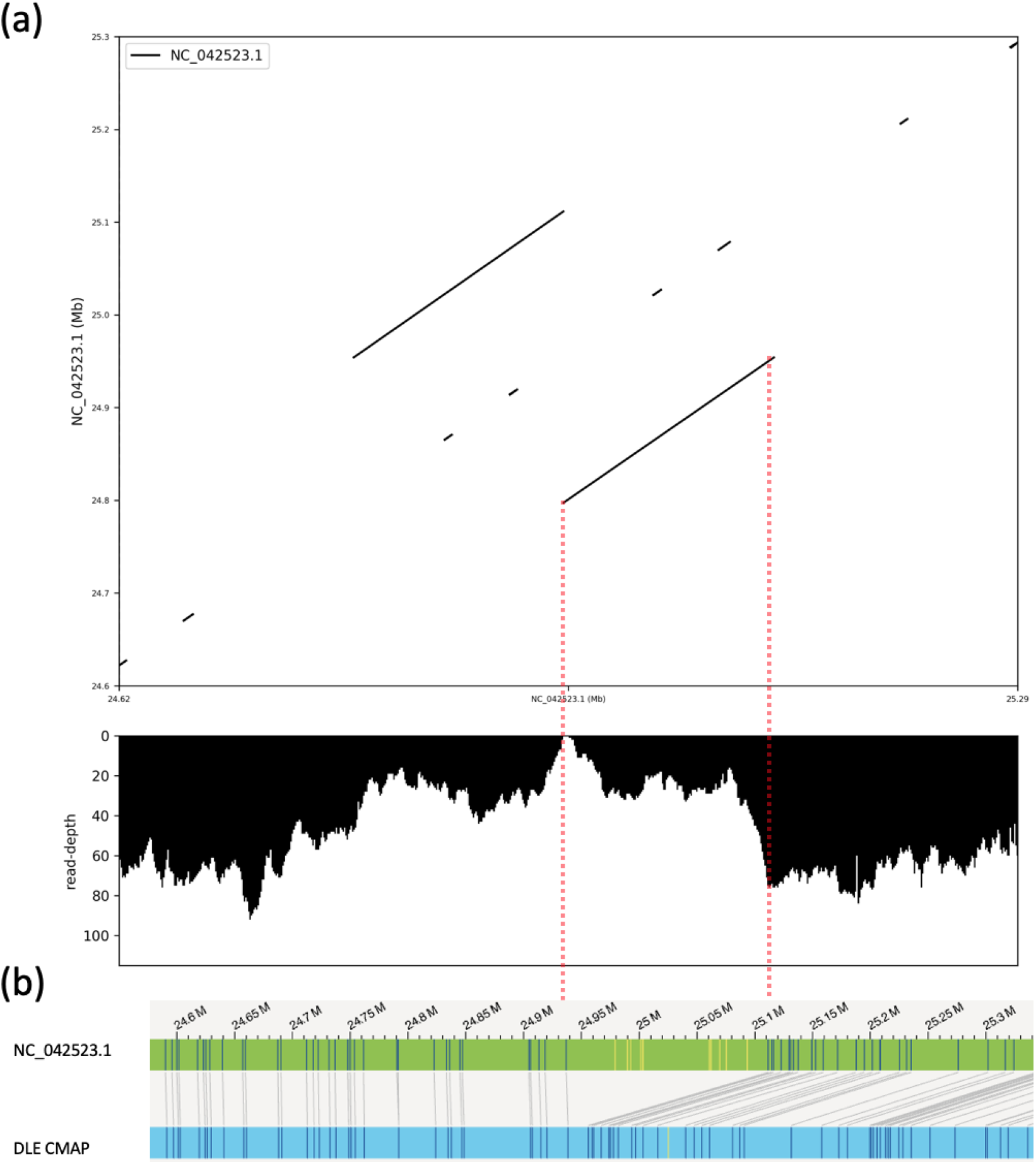
A HAPDUP on fEcheNa1 Chr13 assembly. (**a**) dotplot of self vs self alignment, Chr13 (NC_042523.1) is mapped to itself, and from the PacBio coverage, it is clear the coverage of the matched block drops to about half of the normal coverage. (**b**) the Access view indicates an insertion in the assembly.

Another assembly problem for highly heterozygous assemblies is HAPMIX, in Addtional file 1: Figure S3. It shows an example of a HAPMIX misassembly on Chr5, the two aligned Bionano DLE consensus maps represent each haplotype of the genome, while the draft sequence assembly conflates the haplotypes into one incorrect representation switching between the two.

## Discussion

In this study, we propose a novel method “Asset” to perform misassembly identification based on multiple different types of long-range sequencing data. Through manually checking and comparing with the other tools, we prove that Asset can identify misassemblies accurately, and it discovers much less misassemblies which makes it applicable in the genome curation process.

Here, we think “Asset” has the following features which makes it suitable for assembly evaluation:

Firstly, Asset can evaluate assemblies based on sequencing data. This is the key feature of Asset and makes it valuable to the *de novo* assembly projects such as G10K [8], Bat1K [28]. As more and more newly assembled species are being generated nowadays, reference-based tools are impossible to be applied under most circumstances. Although a reference genome can be available in some rare cases, structure variants between the reference genome and the quality of the reference assembly are still big concerns, using such a reference genome to evaluate the assembly can lead to incorrect outcomes.

Secondly, Asset can identify the misassemblies in long repetitive regions and gap regions more accurately with multiple long-range sequencing data, which is valuable to those genomes containing numerous repetitive structures, such as human (50-70%), mouse (45%).

Thirdly, Asset can generate a limited number of misassemblies, which is helpful during a genome curation process. We have to admit none assemblers are perfect and the genome itself is full of various complex structures, this means even though using a mixed sequencing data to assemble, the final assembly can still involve all kinds of misassemblies. Genome curation is still necessary for producing a high-quality assembly. By using Asset, it can report a suitable number of misassemblies that the genome curators can check easily.

Asset is our first try to integrate supporting information from different sequencing technologies, to discover the structural errors in a draft assembly. This project is far from being finished, we are still checking the misassemblies manually to find out their causalities, more work needs to be done on automatic misassembly classification method and after classification, an automatic program should be applicable to fix those errors as well. Even though Asset can report misassemblies, which can be an indicator for assembly algorithm comparison and can be helpful for generating updated assembly reports including curated N50 or some other metrics, like genome completeness, a comprehensive genome evaluation system will be more useful in the near future. We would consider Asset as our first step to implement such a system. The evaluation system is going to be able to identify, classify, modify and generate updated assembly metrics for our draft or finished assemblies.

As many *de novo* sequencing projects are being launched at the moment, EBP, for example, is chasing its goal to sequence 1.5 million species on earth in the next ten years, which means 150 thousand *de novo* assemblies are coming out on average each year. and we believe it can be imported into the modern assembly pipeline to help the genome curation process and help to evaluate assembly software or pipelines.

## Methods

In this section, we will explain the methods applied on each type of data to find out which sequences or joins are supported, and the method used to merge and filter the suspect misassemblies to get a reliable set of the misassemblies.

### Identification of supported sequences with PacBio data

We first map the raw PacBio data to a given assembly using minimap2 [29] with settings of “-x map-pb”, then we calculate the base level read depth for the assembly with only primary alignment. Then we trim off N bp (default: 300) at the ends of the alignments to capture misassemblies at the boundary of the left and right flanking regions, and calculate read depth for each base. After that, we filter out the low coverage regions where read depth is less than *l* (default: 10) and high coverage regions where reads depth are more than *h* (default: 400). Next, we merged the regions that are less than *b* bp (default: 20) away from each other. Finally, we output the merged regions as PacBio support regions into a bed file. An example is shown in Figure 5a, with the green blocks supported by PacBio data, while the red blocks are not (for illustration purposes the minimum required coverage in this figure is one).

**Figure 5.**
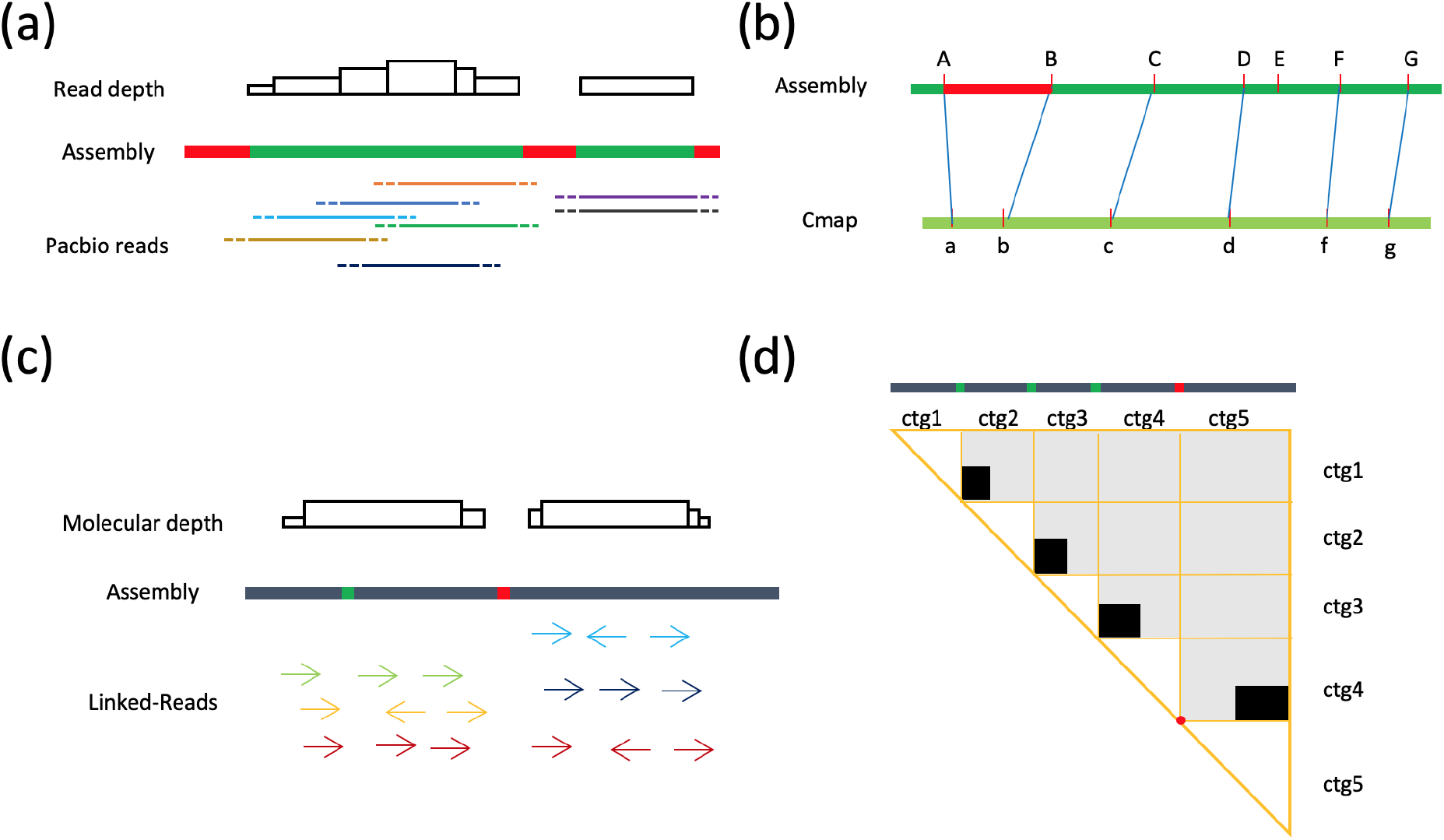
Asset misassembly identification methods. (**a**) supported sequences identification based on PacBio reads. *n* bases are trimmed off at ends (dot lines) of the reads to capture the structural misassemblies at the right and left flanking regions, then coverage for each base is calculated, and sequences whose bases are all above a threshold are output as supported sequences (regions in gray). (**b**) supported regions identification based on the Bionano consensus map. All differences for mapped blocks are collected. An interval for supported regions is estimated based on the differences by using Turkey’s Fence, regions within the interval are treated as supported regions (regions in gray). (**c**) supported joins identification based on linked-reads. The molecules are reconstructed by using only first reads in read pairs, and molecular coverage for each base is computed, joins with molecular coverage in an interval are supported (regions in gray). (**d**) supported joins identification based on Hi-C reads. Contact numbers between all pairs of contigs belonging to the same scaffold are tallied, a join is supported only if its two adjacent contigs should be joined as the order and orientation in the scaffold. As is shown in the subfigure, in this case, ctg1, ctg2, ctg3 and ctg4 are joined correctly, however ctg5 should be flipped and joined with ctg4.

### Identification of supported regions with optical mapping consensus map

Here, we use alignment results between the Bionano consensus map and the assembly to select support regions. We first mapped the Bionano consensus map to the given assembly by RefAligner, the official alignment tool provided by the Bionano company. Then we calculate differences for each mapped block and normalize them by dividing the assembly block length. Next, we used Turkey’s Fence [*Q*1 – 1.5 * (*Q*3 – *Q*1), *Q*3 + 1.5 * (*Q*3 – *Q*1)] which is also applied in [30] to define the thresholds for normalized differences, where *Q*3 and *Q*1 are the third and first quartiles respectively. The mapped blocks having normalized differences within the interval are supported. We output these regions into a bed file. If there are more than one consensus map, Asset will generate supported regions for each of them and merge their supported regions and make one bed file.

As is shown in Figure 5b, alphabets A-G on the assembly and a-g on the consensus map are all digested restriction enzyme sites, and the blue line indicates a match. Regarding the alignment, region “A-B” has a very large divergence with its corresponding region “a-b”, so “A-B” are not supported, while the other blocks are all supported.

### Identification of supported joins with linked reads

In this study, we use 10X linked reads to validate the supported joins. First, we mapped the linked reads to the assembly with defaults settings of “bwa mem” [31]. Then we extract all alignments of the first reads in read pairs, and cluster them by their barcodes, which is usually 16bp in the front of the first read. Here we take all the first reads alignments to guarantee that we have sufficient reads with the same barcode even in the low complexity regions (LCRs) to form complete molecules. Next, we filter out clusters having less than *n* alignments (default: 5). For each cluster, we then ordered its alignments by their mapped targets and loci, merge the alignments that are mapped to the same target and within *l* bp (default: 20kb) away from each other, and calculate average mapping quality *q* for the merged alignments, then drop the merged alignment composed of less than *a* alignments (default: 5) or less than a minimum mapping quality *p* (default: 20). We treat the remaining merged alignments as DNA molecules involved in the sequencing process, the coordinates of the DNA molecule start from the left most read and end at the right most read. We calculate the molecular coverage for each base on the assembly, and calculate average molecular depth *c* for each contig/scaffold, and use to *max*(*r * c, C*) as the minimum molecular coverage, where *C* is a constant and default *r* is set to 0.15. We treat regions larger than the minimum coverage as support regions and output them to a bed file.

As is observed in Figure 5c, the non-gray blocks on the assembly are s between two separate contigs, based on the molecular coverage, the first join which is marked in orange is supported, while the second one is not supported due to its zero molecular coverage.

### Identification of supported joins with Hi-C data

Since Hi-C is not a whole genome sequencing (WGS) technology, we use Hi-C to find supported joins. We first segment the assembly by cutting at the block N’s. Then we map Hi-C data to the segmented assembly using bwa mem with settings of “-SP” which allow the read pairs to be aligned individually. Then we split the contigs into two equal halves and define 5’ end as head (h) of the contig and 3’ end as tail (t), we use a function *F* to represent the link number between contig *i* and contig *j* in a specific link orientation *o,* where *o* can be head to head (hh), head to tail (ht), tail to head (th) and tail to tail (tt),

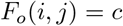

and we also set a function *S* to sum up all links between contig *i* and contig *j*.

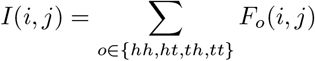

We first initiate *F* with read pairs whose 5’ are mapped unambiguously onto different contigs of the same scaffold without soft or hard clipping and mapping quality is larger than a threshold (default: 10). Then for each contig *i*, we weigh a correlation between *i* and *j* as:

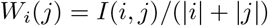

Where |*i*| and |*j*| represent the length of contig *i* and *j* respectively. For contig *i*, we sort *W_i_* in a descending order. Then we iterate all linked contigs *j* in *W_i_*, calculate the contact number *h* and *t* of the head and tail of contig *i* to contig *j*, seek for the first predecessor *p* and successor *s* of contig *i*. We consider *s* a successful successor, if (1) *s* appears after contig *i*, *s* equals *i* + 1, and *t* is larger than *h* or *h* is not significantly larger than *t*; (2) *s + N* appears after contig *i*, *s + N* equals *i* + 1, and *t* is significantly larger than *h*. *N* is configurable by users, the default value is 1. We apply the same way to validate if *p* is a successful predecessor. The details about the algorithm for identifying successful successor of contig *i* are supplied in Additional file 1: Algorithm S1, the same method is applied to check a predecessor.

We can see from Figure 5d, ctg1, ctg2, ctg3, ctg4 are correctly in the natural sequence order and orientation, while ctg4 has a strong signal to join with the tail of ctg5, i.e., ctg5 should be flipped to join with ctg4, so the join between ctg4 and ctg5 are not supported (labeled in red), while other others are supported.

### Misassembly Identification

After we get all the supported sequences and joins, we merge the supported sequences from PacBio and Bionano data, and output the regions that are not supported by any types of the data as our one set of misassemblies, and collect unsupported joins from Bionano, Hi-C and 10X linked-read data as our second set of misassemblies. Then we merge these two sets of misassemblies if they are away within *n*kb (default: 10), after merging, we filter out those misassemblies that are not spanning but are within 1kb to the gaps, they are possible cause by natural drop of PacBio data and missing of Bionano data. The remaining misassemblies are our final suspect misassemblies. All supported sequences and joins from each data type and the final misassemblies are output into bed files, which are easy to be incorporated into genome curation browsers like gEVAL [32].

### Software

The software is implemented in C and is easy to integrate into a modern assembly pipeline. The source code is available at: https://github.com/dfguan/asset.

### Data availability

- CHM13: The assembly is downloaded from https://s3.amazonaws.com/nanopore-human-wgs/chm13/assemblies/chm13.draft_v0.6.fasta.gz. The 215 PacBio RS datasets are available in NCBI under BioProject accession PRJNA269593, 10X, Hi-C and Bionano consensus map are all avaibable from https://github.com/nanopore-wgs-consortium/CHM13.
- Goat: The assembly is available from NCBI Ref-Seq database with accession GCF_001704415.1. The PacBio data we used are all deposited in NCBI Sequence Read Archive (SRA) from SRR3142304-11, SRR3142319-621, and SRR3142754-66, and Hi-C data SRR3773499. The Bionano consensus map is download from https://gembox.cbcb.umd.edu/goat/goat.cmap. The Illumina data used for REAPR are deposited in NCBI from SRR3798850-959.
- fEcheNa1: the fEcheNa1 assembly is deposited in NCBI under BioProject accession PRJEB31992. The reads can be downloaded using the AWS Command Line Interface available at https://vgp.github.io/genomeark/Echeneis_naucrates/

## Author’s contributions

D.G., S.M., and R.D. conceived the Asset project, D.G., S.M., R.D. and G.W. edited the manuscript. D.G. implemented the program and pipeline and ran the experiments. Z.N. and J.W. worked on misassembly validation, and J.W. performed a manual check on the misassemblies for the tested assemblies. Y.Y. generated the HiGlass view for the assemblies.

## Acknowledgements

D.G. and Y.W. were supported by the National Key Research and Development Program of China (Nos: 2017YFC0907503, 2018YFC0910504 and 2017YFC1201201), and D.G. by the China Scholarship Council. S.M. and R.D. were supported by Wellcome grant WT207492, and J.W. and K.H. by Wellcome grant WT206194.

## Additional Files

Additional file 1

Figures S1-S4.

Additional file 2

Table S1. Table of CHM13 suspect misassemblies.

Additional file 3

Table S2. Table of Goat suspect misassemblies.

Additional file 4

Table S3. Table of fEcheNa1 suspect misassemblies.

## References

1. Alkan, C., Sajjadian, S., Eichler, E.E.: Limitations of next-generation genome sequence assembly. Nat. Methods 8(1), 61–65 (2011)

2. Biscotti, M.A., Olmo, E., Heslop-Harrison, J.S.P.: Repetitive DNA in eukaryotic genomes. Chromosome Res. 23(3), 415–420 (2015)

3. Chen, Y.-C., Liu, T., Yu, C.-H., Chiang, T.-Y., Hwang, C.-C.: Effects of GC bias in next-generation-sequencing data on de novo genome assembly. PLoS One 8(4), 62856 (2013)

4. Chin, C.-S., et al.: Phased diploid genome assembly with single-molecule real-time sequencing. Nat. Methods 13(12), 1050–1054 (2016)

5. Myers, E.W.: The fragment assembly string graph. Bioinformatics 21 Suppl 2, 79–85 (2005)

6. Koren, S., Walenz, B.P., Berlin, K., Miller, J.R., Bergman, N.H., Phillippy, A.M.: Canu: scalable and accurate long-read assembly via adaptive k-mer weighting and repeat separation. Genome Res. 27(5), 722–736 (2017)

7. Berlin, K., Koren, S., Chin, C.-S., Drake, J.P., Landolin, J.M., Phillippy, A.M.: Assembling large genomes with single-molecule sequencing and locality-sensitive hashing. Nat. Biotechnol. 33(6), 623–630 (2015)

8. Rhie, M. Arang d Biegler, Iorns, D., Digby, A., Eason, D., Edwards, T., Wilkinson, M., Turner, G., Meyer, A., Kautt, A.F., Franchini, P., William Detrich, H., Svardal, H., Wagner, M., Naylor, G.J.P., Pippel, M., Malinsky, M., Mooney, M., Simbirsky, M., Hannigan, B.T., Pesout, T., Houck, M., Misuraca, A., Kingan, S.B., Hall, R., Kronenberg, Z., Korlach, J., Sović, I., Dunn, C., Ning, Z., Hastie, A., Lee, J., Selvaraj, S., Green, R.E., Putnam, N.H., Ghurye, J., Garrison, E., Sims, Y., Collins, J., Pelan, S., Torrance, J., Tracey, A., Wood, J., Guan, D., London, S.E., Clayton, D.F., Mello, C.V., Friedrich, S.R., Lovell, P.V., Osipova, E., Al-Ajli, F.O., Secomandi, S., Kim, H., Theofanopoulou, C., Zhou, Y., Harris, R.S., Makova, K.D., Medvedev, P., Hoffman, J., Masterson, P., Clark, K., Martin, F., Howe, K., Flicek, P., Walenz, B.P., Kwak, W., Clawson, H., Diekhans, M., Nassar, L., Paten, B., Kraus, R.H.S., Lewin, H., Crawford, A.J., Gilbert, M.T.P., Zhang, G., Venkatesh, B., Murphy, R.W., Koepfli, K.-P., Shapiro, B., Johnson, W.E., Di Palma, F., Margues-Bonet, T., Teeling, E.C., Warnow, T., Graves, J.M., Ryder, O.A., Hausler, D., O’Brien, S.J., Howe, K., Myers, E.W., Durbin, R., Phillippy, A.M., Jarvis, E.D.: Towards complete and error-free genome assemblies of all vertebrate species (2020)

9. Lewin, H.A., Robinson, G.E., Kress, W.J., Baker, W.J., Coddington, J., Crandall, K.A., Durbin, R., Edwards, S.V., Forest, F., Gilbert, M.T.P., Goldstein, M.M., Grigoriev, I.V., Hackett, K.J., Haussler, D., Jarvis, E.D., Johnson, W.E., Patrinos, A., Richards, S., Castilla-Rubio, J.C., van Sluys, M.-A., Soltis, P.S., Xu, X., Yang, H., Zhang, G.: Earth BioGenome project: Sequencing life for the future of life. Proc. Natl. Acad. Sci. U. S. A. 115(17), 4325–4333 (2018)

10. Simão, F.A., et al.: BUSCO: assessing genome assembly and annotation completeness with single-copy orthologs. Bioinformatics 31(19), 3210–3212 (2015)

11. Salzberg, S.L., et al.: GAGE: A critical evaluation of genome assemblies and assembly algorithms. Genome Res. 22(3), 557–567 (2012)

12. Gurevich, A., Saveliev, V., Vyahhi, N., Tesler, G.: QUAST: quality assessment tool for genome assemblies. Bioinformatics 29(8), 1072–1075 (2013)

13. Mikheenko, A., et al.: Versatile genome assembly evaluation with QUAST-LG. Bioinformatics 34(13), 142–150 (2018)

14. Phillippy, A.M., Schatz, M.C., Pop, M.: Genome assembly forensics: finding the elusive mis-assembly. Genome Biol. 9(3), 55 (2008)

15. Hunt, M., et al.: REAPR: a universal tool for genome assembly evaluation. Genome Biol. 14(5), 47 (2013)

16. Jackman, S.D., et al.: Tigmint: Correcting Assembly Errors Using Linked Reads From Large Molecules (2018)

17. Belton, J.-M., et al.: Hi-C: a comprehensive technique to capture the conformation of genomes. Methods 58(3), 268–276 (2012)

18. Howe, K., Wood, J.M.D.: Using optical mapping data for the improvement of vertebrate genome assemblies. Gigascience 4, 10 (2015)

19. Howe, K., Chow, W., Collins, J., Pelan, S., Pointon, D.-L., Sims, Y., Torrance, J., Tracey, A., Wood, J.: Significantly improving the quality of genome assemblies through curation (2020)

20. Yeo, S., Coombe, L., Warren, R.L., Chu, J., Birol, I.: ARCS: scaffolding genome drafts with linked reads. Bioinformatics 34(5), 725–731 (2018)

21. Burton, J.N., Adey, A., Patwardhan, R.P., Qiu, R., Kitzman, J.O., Shendure, J.: Chromosome-scale scaffolding of de novo genome assemblies based on chromatin interactions. Nat. Biotechnol. 31(12), 1119–1125 (2013)

22. Miga, K.H., et al.: Telomere-to-telomere assembly of a complete human X chromosome (2019)

23. Bickhart, D.M., et al.: Single-molecule sequencing and chromatin conformation capture enable de novo reference assembly of the domestic goat genome. Nat. Genet. 49(4), 643–650 (2017)

24. Kerpedjiev, P., et al.: HiGlass: web-based visual exploration and analysis of genome interaction maps. Genome Biol. 19(1), 125 (2018)

25. Dong, Y., et al.: Sequencing and automated whole-genome optical mapping of the genome of a domestic goat (capra hircus). Nat. Biotechnol. 31(2), 135–141 (2013)

26. Ghurye, J., Pop, M., Koren, S., Bickhart, D., Chin, C.-S.: Scaffolding of long read assemblies using long range contact information. BMC Genomics 18(1), 527 (2017)

27. Guan, D., McCarthy, S.A., Wood, J., Howe, K., Wang, Y., Durbin, R.: Identifying and removing haplotypic duplication in primary genome assemblies (2019)

28. Teeling, E.C., Vernes, S.C., Dávalos, L.M., Ray, D.A., Gilbert, M.T.P., Myers, E., Bat1K Consortium: Bat biology, genomes, and the Bat1K project: To generate Chromosome-Level genomes for all living bat species. Annu Rev Anim Biosci 6, 23–46 (2018)

29. Li, H.: Minimap2: pairwise alignment for nucleotide sequences. Bioinformatics 34(18), 3094–3100 (2018)

30. Yuan, Y., Bayer, P.E., Scheben, A., Chan, C.-K.K., Edwards, D.: BioNanoAnalyst: a visualisation tool to assess genome assembly quality using BioNano data. BMC Bioinformatics 18(1), 323 (2017)

31. Li, H.: Aligning sequence reads, clone sequences and assembly contigs with BWA-MEM. PREPRINT 00 (2013)

32. Chow, W., Brugger, K., Caccamo, M., Sealy, I., Torrance, J., Howe, K.: gEVAL – a web-based browser for evaluating genome assemblies. Bioinformatics 32(16), 2508–2510 (2016)

