## Supplementary file for "Genome sequence assembly evaluation using long-range sequencing data"

### Asset Supplementary Note

#### Contents

**1 Supplementary Algorithm** **1**

**2 Supplementary Figures** **1**

#### 1 Supplementary Algorithm

**Algorithm S1:** Hi-C supported join identification algorithm based on N best neighbors and bionimal test

**Input:** sorted $W_i$ as $W$, $N$
**Output:** Bool value for a supported successor $uss$

```
uss ← FALSE
nsp ← 0, nss ← 0, suss ← −1
for k ← 0 to |W| do
    (j, c_hh, c_ht, c_th, c_tt) ← W_k
    t ← c_th + c_tt, h ← c_hh + c_ht
    if j > i ∧ (t > h ∨ bionomial_test(h, 0.5, h + t) ≤ 0.95) ∧ (nss < 1) then
        if j ≠ i + 1 then
            if bionomial_test(t, 0.5, h + t) ≤ 0.95 then
                uss ← TRUE
                for w ← k to k + N ∧ w < |W| do
                    (j, c_hh, c_ht, c_th, c_tt) ← W_w, t ← c_th + c_tt, h ← c_hh + c_ht
                    if j = i + 1 ∧ t > h ∧ bionomial_test(t, 0.5, t + h) > 0.95 then
                        uss ← FALSE
                        break
                    end
                end
            end
        end
        nss ← nss + 1
    end
end
return uss
```

#### 2 Supplementary Figures

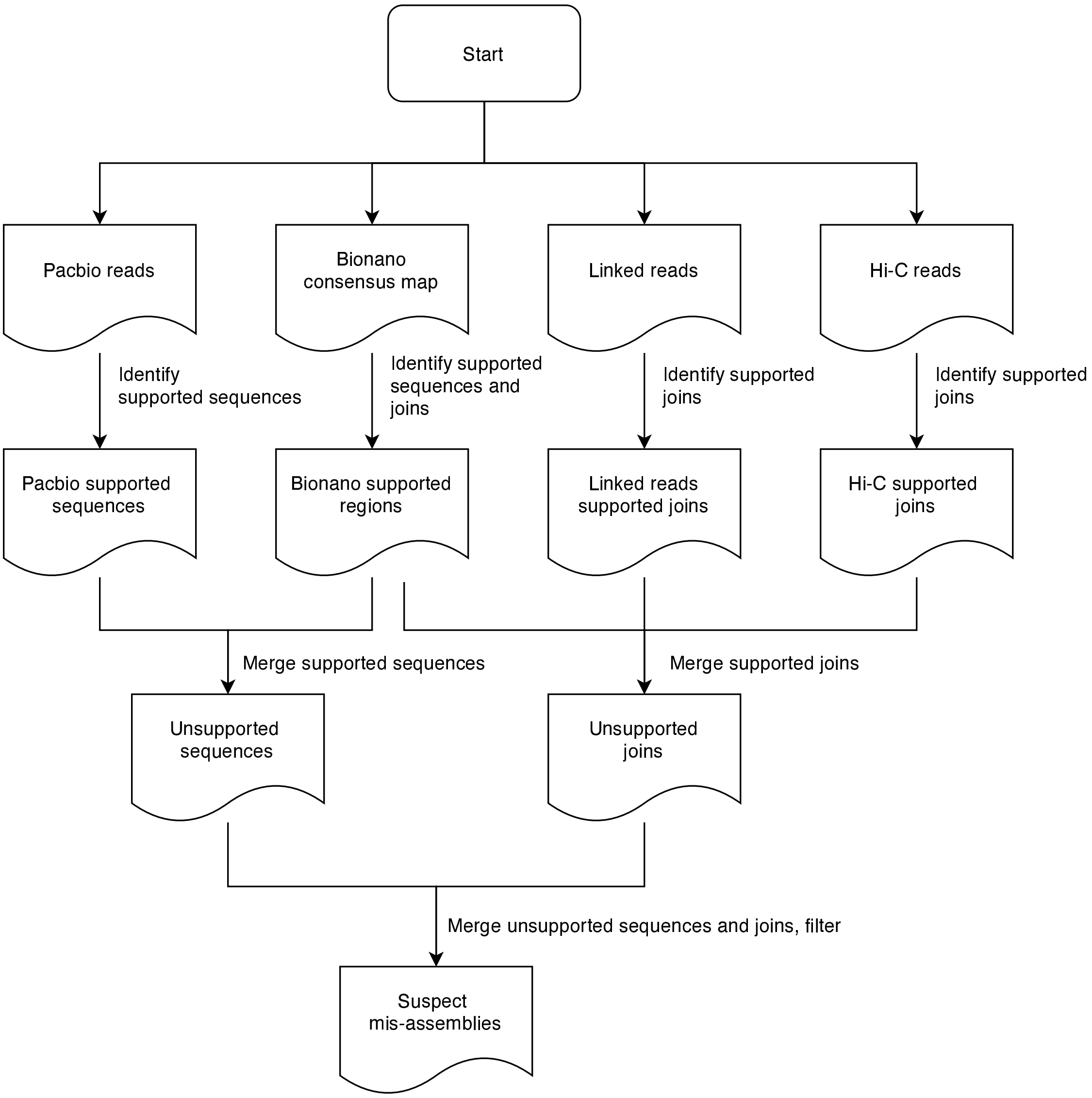


Figure S1: **Asset mis-assembly identification pipeline**. Asset can proceed four types of sequencing data, namely, Pacbio, HiC, Bionano consensus map and linked reads. It takes the inputs, and outputs their supported regions or joins, then it unifies the available supported regions and supported joins, and detect unsupported regions and unsupported joins respectively, and finally asset merges the unsupported regions and joins to make a mis-assembly file.

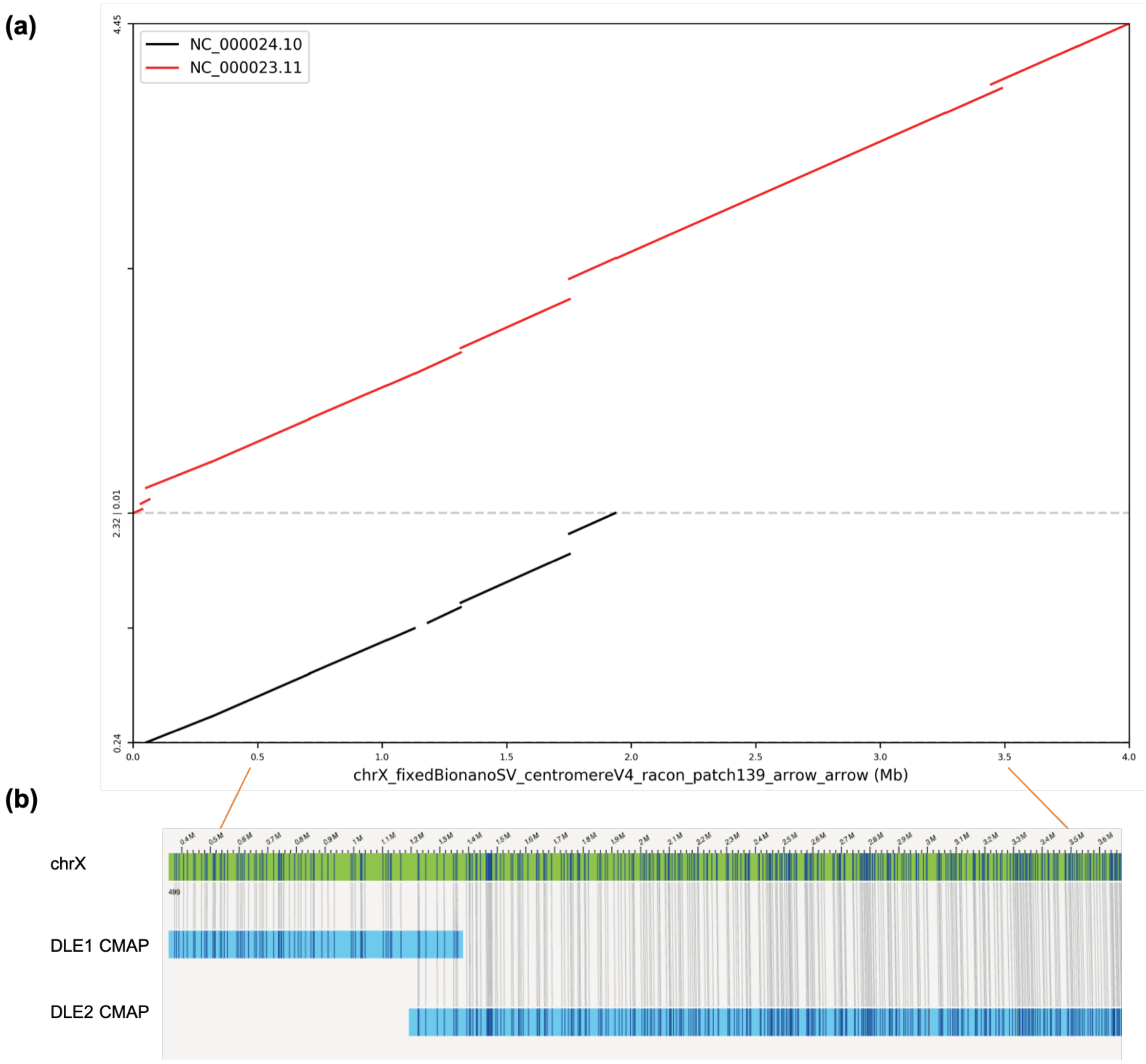


Figure S2: **QUAST-LG identified mis-assemblies on CHM13 assembly**. QUAST-LG identified numbers of translocations. occurring at the beginning of chrX on CHM13 assembly, and based on the dotplot of mapping chrX to GRCh38 assembly, this is due to repeats on chrY assembly.

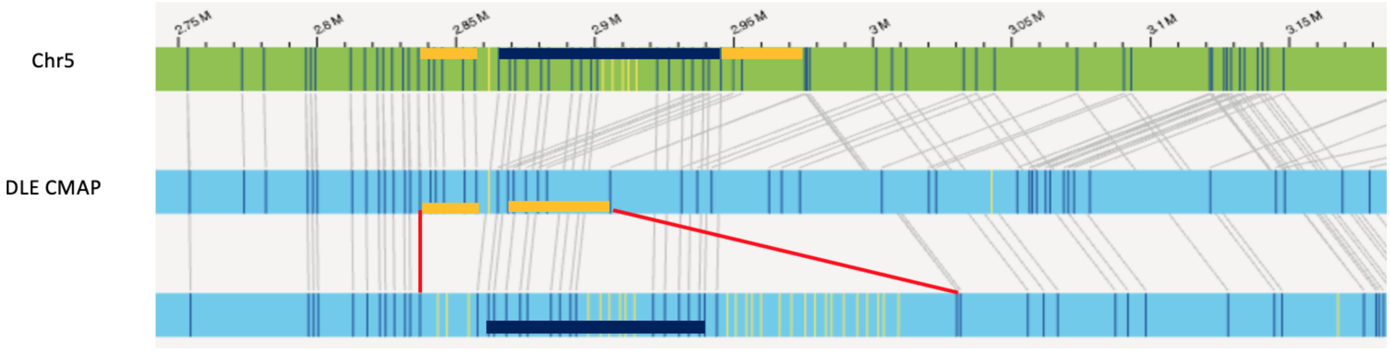


Figure S3: **A HAPMIX on fEcheNa1 chr5 assembly**. IrysView shows a clear divergence between two consensus maps (the red line indicate two different alleles), the assembler assembled the alleles from the first consensus map (yellow thick line), then the second consensus map (blue thick line).
